# how_are_we_stranded_here: Quick determination of RNA-Seq strandedness

**DOI:** 10.1101/2021.03.10.434861

**Authors:** Beth Signal, Tim Kahlke

## Abstract

Quality control checks are the first step in RNA-Sequencing analysis, which enable the identification of common issues that occur in the sequenced reads. Checks for sequence quality, contamination, and complexity are commonplace, and allow users to implement steps downstream which can account for these issues. Strand-specificity of reads is frequently overlooked and is often unavailable even in published data, yet when unknown or incorrectly specified can have detrimental effects on the reproducibility and accuracy of downstream analyses. We present how_are_we_stranded_here, a Python library that helps to quickly infer strandedness of paired-end RNA-Sequencing data.

## BACKGROUND

RNA-Sequencing (RNA-Seq) is the *de-facto* gold standard for the analysis of gene expression on an organism and sample-wide scale — either for the analysis of differential gene expression, transcript structure analysis or identification of novel splice-variants. Common sequencing design of RNA-Seq libraries are either paired-end, where fragments are sequenced from both the 3’ and 5’ end, resulting in two reads per fragment or single-end, where fragments are sequenced from one end only resulting in only one read per fragment (1). Paired-end sequencing libraries result in larger gene transcript coverage, owing to the ability to estimate the distance between the two paired reads and join overlapping reads. This results in improved mapping and subsequently higher accuracy of differential expression analyses, resolution of splice isoforms, and *de-novo* transcriptome assemblies. (2; 3; 4; 5).

Further, library preparation protocols for RNA-Seq can be stranded or unstranded. In unstranded libraries, no information is preserved about the original transcript orientation. In contrast, stranded protocols retain strand information by attaching adapters, or through chemical modification of RNA or the paired cDNA during library preparation (6). Stranded data shows advantages over non-stranded RNA-Seq data such as higher assembly and differential expression accuracy (7; 8). In the case of paired-end data, RNA-Seq eventually results in two fastq files — one for each end of the fragment sequenced. If the data is stranded, we expect all reads from one file to represent the original RNA sequence, and all reads from the other file to represent the complementary cDNA. The two strand-specific layouts can therefore be either fr-stranded, where file 1 contains reads representing the original RNA, or rf-stranded, where file 2 contains reads representing the original RNA. These layouts have varying codes depending on the software used (For reference see (9)). If the data is unstranded, there should be a roughly even and random mix of reads representing the original RNA and reads representing the cDNA in both files.

Downstream RNA-Seq processing pipelines often incorporate information about library design in the workflow, e.g., via a strand-specificity (or strandedness) parameter in RNA assembly and read counting tools. Incorrect use of this parameter can significantly impact the output of RNA-Seq analyses. For example, defining a stranded library as unstranded can result in over 10% false positives and over 6% false negatives in downstream differential expression results (10). Similarly, setting the incorrect strand direction of the RNA-Seq data can result in the loss of >95% of the reads when mapping them back to a reference (11).

RNA-Seq sample strandedness and direction of strandedness is not available as metadata for RNA-sequencing samples in repositories such as The European Nucleotide Archive (ENA) or Sequence Read Archive (SRA), and in the cases where there is a corresponding paper, is often not reported in the methods. From a randomised investigation of 50 ENA “PAIRED END” studies with an associated publication, we found only 56% have strandedness either explicitly stated or mentioned in the methods section for library preparation (Table S1). In addition, we found that the vast majority of papers (94%) do not explicitly state strandedness parameters for downstream software in their methods (Table S1).

Given this lack of reporting, and the impact it can have on downstream analyses we developed how_are_we_stranded_here - a Python library that helps to quickly infer strandedness of paired-end RNA-Seq data. To our knowledge, this is the first stand-alone tool that checks for strandedness. We note that some RNA-Seq tools do issue a warning if the data and stranded parameters do not match (e.g., eXpress (12)) and IRFinder (13)), however these first require full alignment of entire samples to a reference transcriptome or genome, which is time and resource consuming. Similarly, RSeQC’s infer_experiment.py (14) (which we use within how_are_we_stranded_here) requires a genome-aligned BAM file, which again is time and resource consuming to align, making it difficult for users to implement as a standard pre-analysis quality control check.

## RESULTS

### Implementation

how_are_we_stranded_here is written in Python3 and runs a series of commands to determine read orientation. First, a kallisto (15) index of the organisms’ transcriptome is created using transcript fasta sequences, and a GTF which contains the locations and strands for the corresponding transcript sequences. This step is the most time consuming step in the whole process (approx. 6–7 minutes for a human index (15)). As the index remains the same when testing fastq files from the same species, it is saved and can be reused on subsequent tests. Alternatively, pre-built indices can be downloaded from https://github.com/pachterlab/kallisto-transcriptome-indices/releases. Next, input fastq files are sampled to a default of 200,000 reads. These reads are then mapped to the transcriptome, and using kallisto’s-genomebam argument are pseudoaligned into a genome sorted BAM file. Finally, RSeQC’s infer_experiment.py (14) is used to determine the direction of reads from the first and second pairs relative to the mapped transcript, and estimate the number of reads explained by each of the two layouts (FR or RF), and those unable to be explained by either. The output of RSeqQC is then used to calculate the ‘stranded proportion’ — proportion of reads explained by the most common read strand orientation. We expect stranded data to have a stranded proportion of around 1 (i.e. all reads explained by one read direction), and unstranded data to have a stranded proportion of around 0.5 (i.e. half of all reads explained by one read direction, half by the other). If over 90% of read orientation is explained by an FR or RF layout check_strandedness.py outputs the layout as the most likely sequencing layout. Similarly, if neither of the two layouts can be explained by more than 60% of the reads the data is reported as unstranded.

### Testing on simulated reads

We first tested how_are_we_stranded_here on simulated samples across three species — human (Homo sapiens), yeast (Saccharomyces cerevisiae), and thale cress (Arabidopsis thaliana). Using lower numbers of reads resulted in greater variation in percent stranded reads for samples with non-strand-specific reads (one-sided F-test; p<0.05 for all comparisons in read counts different by one order of magnitude; Table S2) in all species (Fig. 1, S1 A). We found that at least 200,000 reads were required to call percent stranded within 0.5% (3σ), and therefore recommend use of 200,000 reads — which is also the default setting for RSeQC. Further, we found very little difference between reported strandedness percentages and known percentages when using 200,000 reads (mean average error < 0.1%), confirming that this method indeed accurately reports stranded percentages in simulated data (Fig. S1 B).

**Figure 1.**
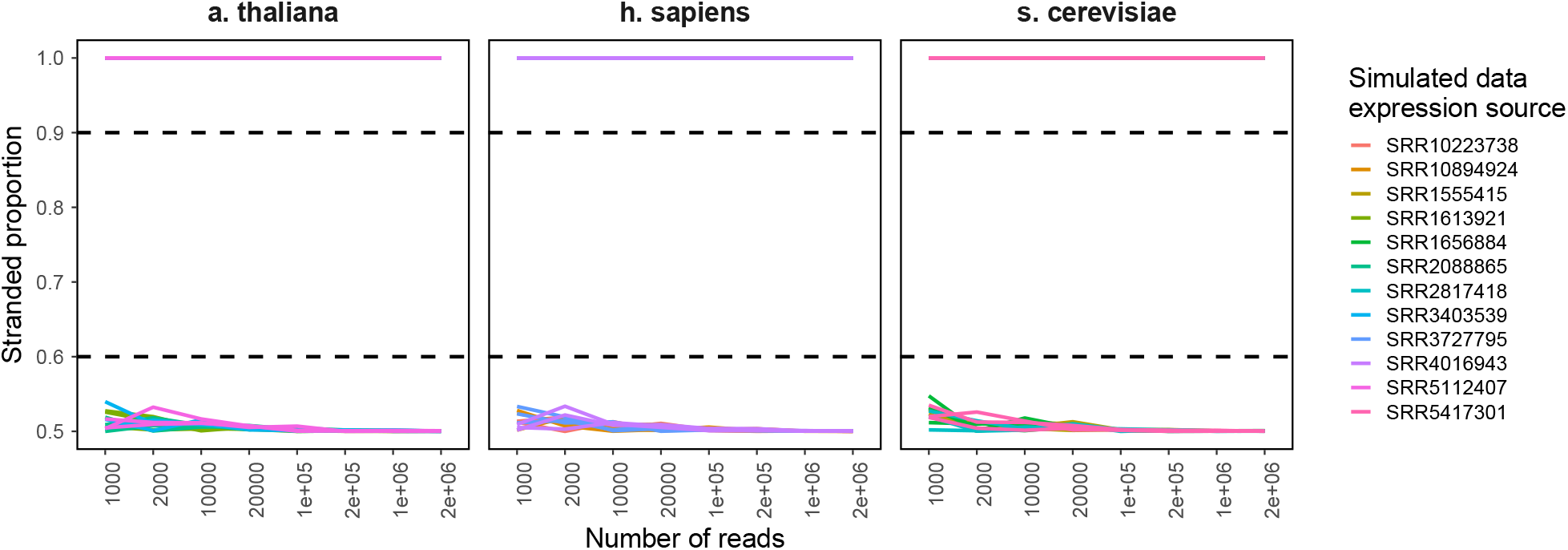
Strandedness proportions in simulated RNA-Seq data. Four biological samples for each species were used to generate three simulated replicates each using polyester (16) at varying read numbers with either strand-specific or non-specific reads. All samples show the correct strandedness, with the strandedness proportion was below 0.6 (unstranded) or above 0.9 (stranded; dashed lines). how_are_we_stranded_here was run using the full Ensembl cDNA annotation for each species.

We then tested the speed of running how_are_we_stranded_here on simulated data using 200,000 reads. Each run took less than 45 seconds for human, 10 seconds for yeast, and 20 seconds for thale cress on a 2020 M1 Macbook pro with 16GB RAM (Table S3).

### Testing on ENA studies

We next tested how_are_we_stranded_here on samples from 60 studies across the same three species as above. The majority of samples showed either a stranded proportion of greater than 0.9 (stranded) or less than 0.6 (unstranded), when removing data points where less than 10% of sampled reads aligned to the transcriptome (Fig. 2). Reported proportions were similar with increasing reads sampled, with a median difference of 0.5 percentage points when using 1000 reads compared to 2M reads (Table S4). As noted before we recommend using 200,000 reads for alignment, due to its higher similarity in reported proportion and decreased alignment time.

**Figure 2.**
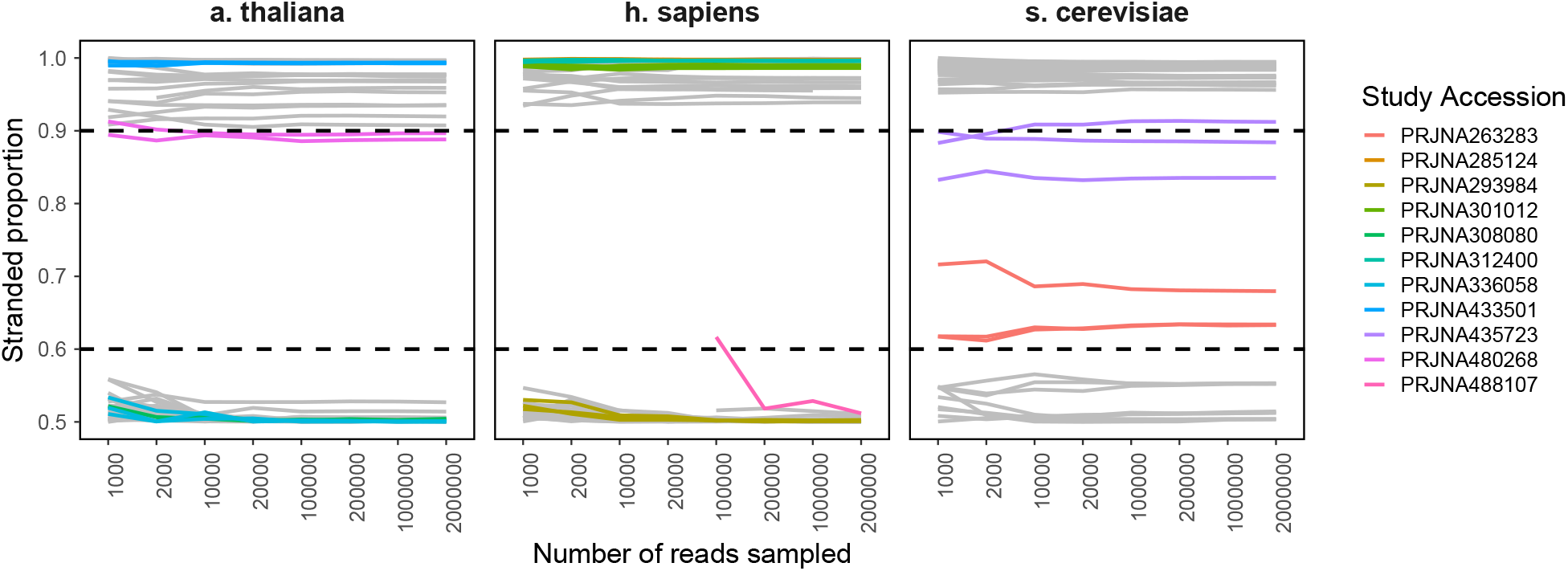
Strandedness proportions in RNA-Seq data. Strandedness proportions were evaluated for 20 studies for each h. sapiens, s. cerevisiae, and a. thaliana using how_are_we_stranded_here and varying the number of input reads sampled. Results are shown where the proportion of reads psuedoaligned is at least 0.1. Studies for which the strandedness proportion was between 0.6 and 0.9 (dashed lines), and those which do not match the reported strandedness are highlighted. how_are_we_stranded_here was run using the full Ensembl cDNA annotation for each species.

While the majority of samples matched their reported strand-specificity, there were seven which did not (Fig. 2, Table S5). We appreciate that this may have been due to inaccuracy in reporting library preparation kit names, as methods such as Illumina’s TruSeq library preparation kits have multiple variants with similar names. For example, the TruSeq Stranded mRNA kit (20020594), and the TruSeq RNA Library Prep Kit v2 (RS-122-2001) could both be referred to as a Illumina TruSeq library preparation kit, but produce stranded and unstranded reads respectively. This may have also been due to unclear communication of methods in general and mix-ups in methods reporting, again highlighting the importance of the use of how_are_we_stranded_here as a quality check on raw RNA-Seq data.

Two yeast studies showed a stranded proportion between 0.6 and 0.9 (Fig. 2; PRNJA263283 and PRNJA435723). Both of these studies were reported as stranded (Table S5). To investigate why the proportion of stranded reads was lower than expected, we investigated the proportion of reads that aligned to specific features. Furthermore, by aligning the data to other species we tried to exclude sample contamination or sample mix ups. None of the samples showed a significant amount of reads that mapped to other species (Table S6), suggesting that sample contamination from another species did not cause the ambiguous results. We then used STAR to align reads to the s. cerevisiae genome, and HTSeq-count to count reads aligning to genes and intergenic regions (see Methods). We found that SRR6767450 contained a greater number of reads aligning to small RNAs (Fig. S2 A), suggesting incomplete removal of small RNAs. Similarly, SRR1605749 displayed the greatest number of reads aligning to intergenic regions, which could be explained by RNA contamination with genomic DNA (Fig. S2 A). We further explored this by running how_are_we_stranded_here on intergenic regions only (see Methods), which showed a stranded proportion of 50%, consistent with the expectation that genomic DNA shows no strand bias (Fig. S2 B).

We further investigated if quality and adapter trimming may have any effect on strandedness proportion, and found that although several samples had lower quality reads and adapters, trimming had very little effect on strandedness (Fig. S3 A). In addition, screening for contaminating reads and removing multimappers had also had little effect on strandedness proportions (Fig. S3 B).

## DISCUSSION AND CONCLUSIONS

Here we presented how_are_we_stranded_here, a tool for quick determination of strandedness in RNA-Seq data. Strandedness checks should be performed prior to any downstream processing of RNA-Seq data — for which many tools require a strandedness parameter which if incorrectly assigned can produce inaccurate data. We recommend strandedness to be checked on at least three samples in the first instance, and on further samples when strandedness is contradictory in these samples, or the strandedness proportion is between 0.6–0.9. This quality control check allows users to confirm strandedness of data when the library preparation method was known, and estimate strandedness when the method is unclear. We show that it may also be indicative of sample contamination, and can warrant further quality control checks. While library preparation (and the strandedness parameters in downstream processing) should be detailed in the methods of published RNA-Seq data, as this is vital to being able to reproduce results, we found that over a third of publications do not provide this detail. As such, how_are_we_stranded_here allows users to easily find the correct strandedness parameter for RNA-Seq datasets which is crucial for reproducibility of published results.

## METHODS

### Screening for RNA strandedness in ENA RNA-Seq studies

All read runs matching the rules library_strategy=“RNA-Seq”, library _source=“TRANSCRIPT()MIC”, and library _layout=“PAIRED” were retrieved from ENA (17). The runs were then randomly ordered, and the first 50 with an associated publication were selected for manual screening of methods. Methods were searched for the library preparation details or kit name, and any mention of “strand” or “direction”. In cases where only the library preparation kit was named, we searched through the company’s product descriptions to discern if the kit was capable of producing strand specific data. Further, if none of these details were found in the methods, we searched the main text for any mention of “strand” or “direction”. Further, we searched the computational methods — either within the publication, supplementary data, or in a code repository — for any strand-specific parameters.

### Data set selection for testing

All read runs matching the following rules were retrieved from ENA for each human (h. sapiens; taxid 9606), yeast (s. cerevisiae; taxid 4932) and thale cress (a. thaliana; taxid 3702): instrument_platform = “ILLUMINA”, library_strategy=“RNA-Seq”, library_layout=“PAIRED”, library_source=“TRANSCRIPTOMIC”, library_selection=“cDNA”. These runs were then filtered for those sequenced on an Illumina HiSeq 2000, 2500, 3000, or 4000, between 10 and 30 samples per study, and randomly reordered. Studies were searched for any publications that were associated with the data, and only retained if strandedness was stated or able to be inferred by the library preparation methods (see above). The first 20 studies matching these requirements were used for analyses for each species.

### Generation of simulated RNA-Seq reads

Reads were simulated using polyester (16). Kallisto 0.46.1 (15) with the corresponding Ensembl 100 transcript annotation (18) was used to count reads for each transcript from an original sample (see above) for four samples per species, two with strand-specific reads and two with non-specific reads. These read counts were then used as the basis to generate three simulated samples each with relative transcript abundances as would be expected in real samples. Reads were then simulated for each of these samples (12 per species), as both strand-specific, and non-specific in multiple numbers of reads (1000-2M).

### Testing of how are we stranded here

For each study the first three samples were taken to profile for strandedness. A kallisto (15) index was generated for each species, using the Ensembl 100 transcript annotation (18). Each sample for the ‘real’ dataset was then run through how_are_we_stranded_here, with varying numbers of reads — from 1000 to 2 million. For the simulated datasets, samples were generated for varying numbers of reads in both strand-specific and non-specific formats. Simulated mixed samples were generated by combining a random subset of reads from the same strand-specific and non-specific samples at varying ratios to give a total of 200000 reads. Simulated datasets were similarly run through how_are_we_stranded_here, using the same number of reads as was simulated (i.e., not sub-setting the same sample). The actual number of reads that were pseudoaligned by kallisto was checked from the kallisto output log. The “stranded proportion” was calculated by taking the maximum proportion of reads explained by RF or FR.

### Quality checks

All steps were performed on samples of 200000 reads from each fastq file, to match the number of reads used for how_are_we_stranded_here. FastQC 0.11.5 (Babraham Bioinformatics) was used to assess the quality of reads, and Trimgalore! 0.6.0 (Babraham Bioinformatics) in paired end mode to quality trim reads and remove adapter sequences. We used FastQ Screen 0.14.0 (Babraham Bioinformatics) to screen for contaminants in trimmed fastq files, with the default databases downloaded by “1’as(qscreen-get_genomes”. Reads pairs which both mapped only to the correct genome were extracted for each sample using the-tag flag in FastQ Screen, and a custom R script (see below). We then ran how_are_we_stranded_here on the trimmed, and the exclusively correctly-mapping fastq files. For genome alignment with STAR 2.7.0e (19), the first sample for each study was aligned to the Ensembl 100 s. cerevisiae genome using default settings. The index was prepared using default settings, the s. cerevisiae genome fasta, and the s. cerevisiae Ensembl 100 GTF. An ‘intergenic’ GTF annotation was generated by creating features covering fully intergenic regions (i.e. not overlapping annotated genes on either strand) and then subdividing these into smaller features using exomeCopy::subdivideGRanges (20) and a subsize of 1000. All intergenic regions were assigned the “+” strand, to allow for calculation of stranded proportions. For read counting, HTSeq-count 0.12.4(21) was used with a GTF annotation containing both the reference Ensembl gene annotation and the intergenic annotation. Read counts were then summed for each gene biotype — including ‘intergenic’. To find the stranded proportion of intergenic reads, how_are_we_stranded_here was run 200,000 reads and using the intergenic annotation instead of the reference annotation.

## AVAILABILITY OF DATA AND MATERIALS

### Code availability

The how_are_we_stranded_here python package is available on GitHub in the repository https://github.com/betsig/how_are_we_stranded_here under a MIT license. All scripts used in the analyses performed in this manuscript are available at https://github.com/betsig/strandedness_testing_scripts.

### Data availability

Public datasets were downloaded from the European Nucleotide Archive. Sample identifiers are listed in Table S4.

