## Supplementary Figures for "how_are_we_stranded_here: Quick determination of RNA-Seq strandedness"

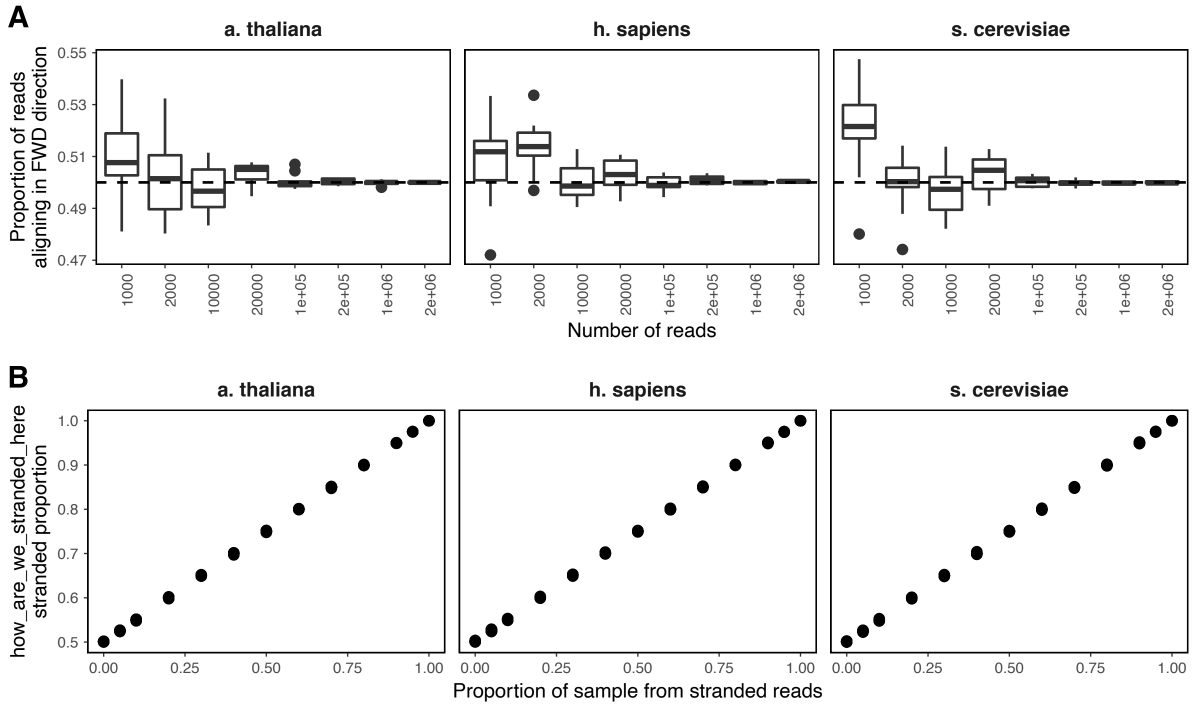


**Figure S1: Strandedness proportions in simulated data.** A. Range of reported forward strand-aligning proportions for non-strand-specific data at varying numbers of reads. Lower numbers of reads show greater variability in reported proportions. B. Reported stranded proportions in simulated data with known stranded to unstranded read proportions. All samples in B. contained a total of 200,000 reads.

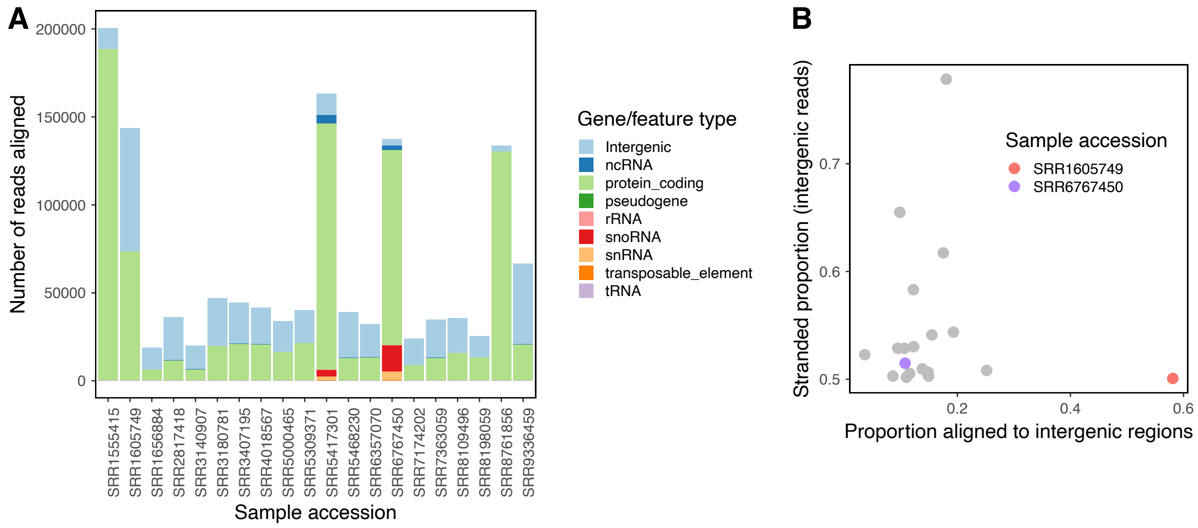


**Figure S2: Reads mapping to intergenic regions in s. cerevisiae samples.** A. Number of reads aligned to each gene biotype or feature using STAR and RSEM. Biotypes were obtained from the Ensembl GTF annotation. Intergenic regions were defined as in Methods. Reads not aligned to any feature are not shown. B. ho_are_we_stranded_here stranded proportion in reads aligning to intergenic regions. Two samples with unclear strandedness from initial checks are highlighted.

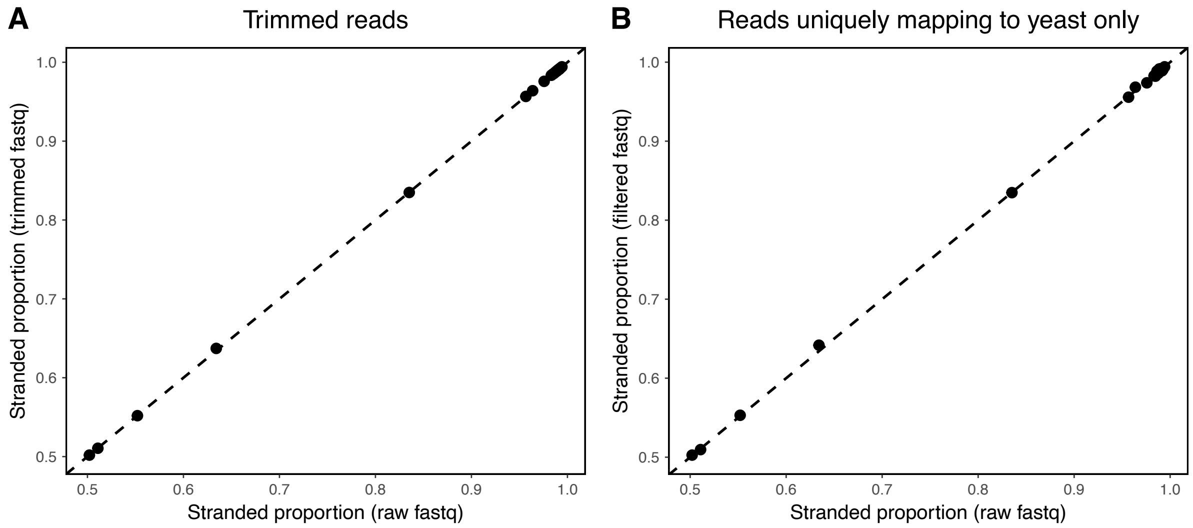


**Figure S3: Strandedness proportions are not altered by trimming or filtering reads.** A. Stranded proportions of yeast samples in raw fastq files and adapter filtered/quality trimmed fastq files. B. Stranded proportions of yeast samples in raw fastq files and fastq files containing reads that uniquely map to yeast. Uniquely mapping reads were obtained using FastQ Screen and extracting pairs which had one hit to the yeast genome.
